# Exploring the Neural Basis and Validity of Ordinal Emotion Representation through EEG

**DOI:** 10.1101/2024.11.21.624627

**Authors:** Xuyang Chen, Xin Xu, Dan Zhang, Xinke Shen

## Abstract

Accurately measuring emotion is a major challenge in advancing the understanding of human emotion and developing emotional artificial intelligence. In many existing studies, participants’ emotional ratings in interval scales are considered the true reflection of their emotional experiences. However, recent research suggests that ordinal annotations of emotions can more accurately capture the emotional expression process, providing a potential method for more precise emotion measurement. However, our understanding of the characteristics and validity of this new form of emotion representation is still relatively lacking. In particular, there is a lack of research using neural signals to explore the validity and neural basis of ordinal emotion representation. In this study, we used a video-elicited electroencephalogram (EEG) dataset (n = 123) to identify the neural basis of ordinal emotion representation and demonstrate its validity from a neural perspective. Furthermore, we explored various characteristics of ordinal emotion representation, showing how it is superior to the interval form. First, we conducted inter-situation representational similarity analysis (RSA) and inter-subject RSA to test the degree to which ordinal representation captures both group commonalities and individual differences of emotion. Next, we investigated the characteristics of ordinal representation under different combinations of emotion items, including uni-variate and multivariate emotions, positive and negative emotions. Our results show that both group commonalities and inter-subject variations in EEG features are better explained by ordinal emotion representations than by interval ones. Multivariate ordinal representations showed better inter-subject reliability and higher representational similarity with EEG features compared to uni-variate counterparts, highlighting the co-occurrence nature of human emotions. Compared to negative emotions, ordinal representation showed greater improvements for positive emotions, suggesting that the complexity of positive emotions is well captured by ordinal representations. Taken together, these findings demonstrate that multivariate ordinal emotion ratings provide a more accurate measure of real emotional experience, which is crucial for enabling machines to precisely understand and express human emotions.

## 1 Introduction

Grasping human emotions and measuring them with precision remains a core challenge in psychological research and the advancement of emotional artificial intelligence (AI). Emotional AI seeks to empower machines to recognize, interpret, and respond to human emotions, showing great potential for enhancing human-computer interactions, advancing mental health diagnostics, and developing more empathetic technologies. Achieving accurate emotion measurement is critical for advancing these areas, as it ensures emotional AI can effectively comprehend and engage with human emotional states.

However, current emotion measurement techniques have significant limitations, mainly stemming from the widespread use of interval-based approaches^[1]^. Interval data poses challenges for capturing subjective constructs like emotions, as it may inaccurately reflect a person’s actual emotional state. Such misrepresentation occurs because the externalization of emotion experience is inherently relative rather than absolute^[2,3]^. This makes the absolute differences in interval scores unstable and noisy^[2,4]^, thus not suitable for describing emotions. Additionally, interval measurements often exhibit low reliability, especially in dynamic contexts where emotions fluctuate over time^[3]^. These limitations underscore the need for more precise methods of emotion annotation.

Recent research has proposed that ordinal representations may provide a more accurate reflection of emotional experiences^[3]^. Unlike interval ratings, ordinal scales are more aligned with how humans naturally perceive and express emotions, which are inherently comparative. Empirical evidence highlights several benefits of ordinal annotations: 1) Greater inter-rater reliability: Ordinal scales reduce subjective biases and inconsistencies commonly found with interval ratings, leading to more consistent results across different raters^[3,5]^. 2) Stability and robustness: Ordinal measures capture the rank order of emotional intensity without assuming exact differences between emotions, making them less vulnerable to distortions caused by extreme ratings^[6,7]^. 3) Better alignment with human perception theories: Ordinal data more accurately reflects the comparative nature of emotional experiences, avoiding the pitfalls of misrepresenting emotional states through fixed interval scales^[3]^. These benefits suggest that ordinal emotion representations may better capture the structure of emotional experiences and provide a promising alternative to traditional interval measurement methods^[5]^.

However, the validity of ordinal measurements of emotions is not fully investigated. Existing research on the validity of ordinal forms of emotion has been limited to peripheral physiological (e.g., skin conductance^[8–10]^, heart rate^[10]^) and non-physiological signals such as speech^[8]^. Currently, there is a lack of validation based on neural signals, testing whether ordinal forms of emotion better corresponds to the neural activity in human brain. Validating the ordinal form of emotion using neural signals can demonstrate its neurophysiological substrates. Uncovering the neural basis of different emotion categories is crucial for understanding the underlying process of emotion externalization. Besides, the current understanding of the characteristics of this new form of emotion representation is still lacking — how is ordinal form superior, and what specific aspects of emotions it can better describe, remain largely unknown.

To address the above issues, we used a video-elicited emotional electroencephalogram (EEG) dataset with 123 subjects to conduct validation based on neural signals and identify the neural representations of ordinal emotions. Next, we explored the performance of ordinal emotions in terms of group commonalities, individual differences, uni-variate/multivariate emotions, and positive/negative emotions. To be specific, first, recent studies have proposed inter-subject representational similarity analysis (RSA) and inter-situation RSA as powerful methods for studying individual differences and group commonalities in emotions^[11,12]^. These methods can also be used to reveal how well the ordinal form describes individual differences and group commonalities in emotional experiences, in a way that aligns with the neural signal. Second, analyses under different combinations of emotional items can demonstrate the characteristics of the ordinal emotions on multiple aspects. For example, dividing emotions into positive and negative categories can show the relationship between these two primary emotional groups and the ordinal form; viewing emotions as multidimensional vectors can capture the complexity and co-occurrence of emotional experiences, allowing for a more nuanced characterization of emotions^[13]^.

## 2 Methods

### 2.1 Data Acquisition and Processing

#### 2.1.1 The FACED Dataset

To validate the proposed algorithm, we used data from the Finer-grained Affective Computing EEG Dataset (FACED)^[14]^. The FACED dataset aims to provide fine-grained emotional computing data, with a balanced classification of both positive and negative emotional aspects.

This dataset contains 32-channel EEG recordings from 123 participants, each of whom watched 28 emotionally evocative video clips categorized into nine distinct emotions: amusement, inspiration, joy, tenderness, anger, fear, disgust, sadness, and neutral. Each negative and positive emotion category included three video clips, while the neutral category featured four. The clips had an average length of about 66 seconds, ranging from 34 to 129 seconds. After viewing each clip, participants were asked to report their subjective emotional experiences across 12 items, covering eight specific emotions (anger, fear, disgust, sadness, amusement, inspiration, joy, tenderness) and four broader dimensions (valence, arousal, liking, familiarity). Ratings were provided on a continuous scale from 0 to 7, where 0 represented “very negative” for valence and “not at all” for the other dimensions, while 7 represented “very positive” or “very much.” The analysis focused on multivariate ratings across the eight specific emotion categories.

#### 2.1.2 EEG Data Preprocessing

The EEG signals were downsampled to 125 Hz and bandpass filtered between 0.5 to 40 Hz using a fourth-order zero-phase-shift Butterworth filter. Noisy channels were identified manually by assessing variance and outlier values, with an average of 0.06±0.26 noisy channels per trial. These noisy channels were interpolated using the average of three neighboring channels. Independent component analysis (ICA) was applied to eliminate ocular and muscle artifacts, with 2-4 components being removed per participant. The data were then re-referenced to the average of the two mastoid channels. Preprocessing steps utilized Noisetools^[15]^ and Fieldtrip toolboxes^[16]^.

### 2.2 The Computation of Distance and Ordinal Representations

The core idea behind ordinal representation is that individuals evaluate emotions relatively. Thus, we transformed the emotional annotations for each participant individually. To extract ordinal information from the interval annotations (Fig. 1a), we first calculated the relative differences within each dimension to derive a distance representation (Fig. 1b). Next, the magnitude of the differences, which is irrelevant in ordinal representation, was discarded, leaving only the sign (indicating the greater/lesser relationship in ordinal representation, Fig. 1c). The specific computational process is outlined below.

**Fig. 1.**
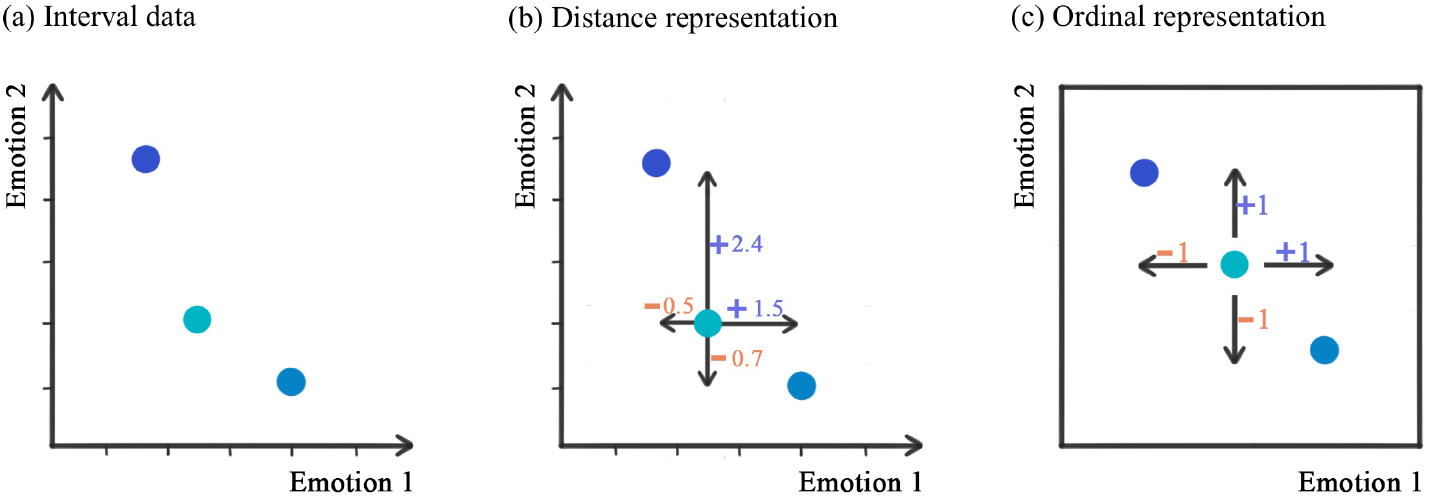
General idea of extracting ordinal representation from interval annotations. (a) the original annotation data in interval scale. (b) Considering the relative positions on each emotion dimension. (c) Only preserving the rank information yields the ordinal representation.

#### 2.2.1 Distance Representation

For a participant’s annotations on *E* emotions across all *N* situations, *Y*_1_, *Y*_2_, …, *Y*_*n*_, where *Y*_*i*_ ∈ ℝ^*E*^, all annotations forms a network with *N* nodes and *N ×* (*N* − 1)*/*2 edges. Rather than expressing the emotion of each trial in its original emotional space, we represent emotions based on their relative positions within the network. In this network, the position of each node is determined solely by its relative relationship to other nodes, making it independent of absolute coordinates. These relative positions are described through a set of distance vectors, and this representation is referred to as the distance representation. Specifically, for *Y*_*i*_, the distance representation consists of a set of distance vectors {*Dist*_*ij*_|*j* = 1, 2, …, *i*, …, *N* }, where:

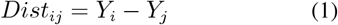

The vector *Dist*_*ij*_ ∈ ℝ^*E*^ represents the distance between node *Y*_*i*_ and *Y*_*j*_. By concatenating the one-dimensional distance vectors, we generate the distance representation matrix *D*_*i*_ ∈ ℝ^*E×N*^ .

#### 2.2.2 Ordinal Representation

In ordinal measurement, the specific numerical differences between two annotations hold no intrinsic meaning. Therefore, we convert the distances into ordinal categories—greater than, less than, or equal to. This transformation shifts the focus from the actual magnitude of differences to the relative ordering of the nodes, capturing the ranking relationships instead. This process is represented as follows:

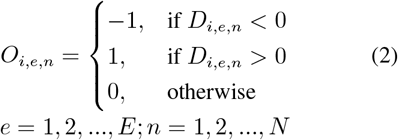

Where the values 1, 0, and -1 are used to represent “greater than,” “equal to,” and “less than” in ordinal measurement. The transformed matrix *O*_*i*_ ∈ ℝ^*E×N*^ captures only the size relationship information from the distance representation, while discarding the numerical values. This ordinal representation retains only the relative ranks of the annotations, making it equivalent to ordinal measurement.

### 2.3 Inter-rater Reliability

Inter-rater reliability evaluates the level of agreement among a group of raters^[17]^ and is widely used to assess the reliability of emotion annotations, and also an important indicator of whether an emotion measure is reasonable. Previous research has shown that ordinal emotions have higher inter-rater reliability than interval emotions. Here, we further examine the reliability improvements provided by ordinal representations, considering both univariate/multivariate and positive/negative emotions. We assess inter-rater reliability by measuring the consistency of the inter-situation similarity matrix across participants. A higher consistency indicates that participants rate emotions in a similar manner across different situations, as compared to how other participants rate them. In this study, we use the four-dimensional vector composed of “joy,” “tenderness,” “inspiration,” and “amusement” to represent positive emotions, and the four-dimensional vector composed of “anger,” “disgust,” “fear,” and “sadness” to represent negative emotions. On the other hand, an eight-dimensional vector composed of all eight emotion items is used to represent multivariate emotions, while each single emotion item represents uni-variate emotions. We then calculate each participant’s inter-situation emotion similarity matrix based on both ordinal and non-ordinal emotion representations, and compute the consistency across participants.

#### 2.3.1 Inter-situation Similarity Matrix of Ratings for Each Subject

First, for each participant, we calculated the inter-situation similarity matrix of their ratings. Here, we denote 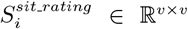 as the inter-situation similarity matrix of emotion annotations for participant *i*. It is computed based on the annotations across all situations for that participant, *R*_*i*_. When using multivariate emotion, *R*_*i*_ ∈ ℝ^*v×E*^ includes subject *i*’s ratings of *v* situations (i.e., elicitation videos) on all *E* emotion items; When using uni-variate emotion, 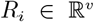 includes subject *i*’s ratings of *v* situations on one specific emotion item; When using positive emotion, 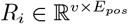 includes subject *i*’s ratings of *v* situations on *E*_*pos*_ positive emotion items; When using negative emotion, 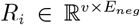 includes subject *i*’s ratings of *v* situations on *E*_*neg*_ negative emotion items.

For any two situations *v*_1_ and *v*_2_, calculate the Euclidean distance between 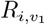,: and 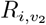, resulting in the distance matrix 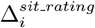.

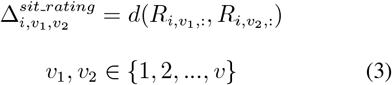

Where *d*(*x, y*) is the *L*_2_ distance between *x* and *y*. 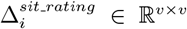 represents the subject i’s ratings dissimilarity across v situations.Then, the dissimilarity matrix was converted into similarity matrix 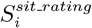 by normalizing it and subtracting it from an all-ones matrix.

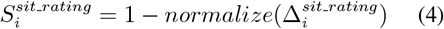

#### 2.3.2 Inter-rater Consistency of Inter-situation Similarity Matrices

After obtaining the inter-situation similarity matrix for each participant’s ratings, which is 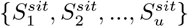,the next step is to compute their inter-rater consistency. We enumerate all possible pairings between all participants and calculate the Pearson’s correlation of their similarity matrices. The resulting distribution of all *u ×* (*u* − 1)*/*2 r-values demonstrates the inter-rater consistency of the inter-situation similarity matrices. By definition, this reflects the inter-rater reliability of the emotion ratings.

### 2.4 Representational Similarity Analysis

Representational Similarity Analysis (RSA) is a commonly used multivariate analysis method in cognitive neuroscience, designed to quantify and compare information representations in the brain or computational models. RSA evaluates representations in a brain region or computational model by constructing a distance matrix of response patterns, highlighting which differences between stimuli are accentuated and which are minimized^[18]^. In this study, we perform two types of RSA (Fig. 2): inter-situation RSA and inter-subject RSA. Inter-situation RSA involves comparing the similarity patterns of neural signals across different situations with the similarity patterns of emotion ratings (where “situations” refer to the emotion-eliciting materials). This analysis aims to test how well the situation-varying patterns in emotion ratings shared by all individuals are aligned with those of the neural signals. The inter-situation RSA was indicative of “shared” emotion ratings as it was conducted based on the group-averaged neural similarity patterns across situations and the group-average rating similarity patterns. The variation in group-averaged similarity matrices reflected the “shared” response across participants. Inter-subject RSA, on the other hand, compares the similarity patterns of neural signals across participants with the similarity of emotion ratings across those participants. This analysis aims to test how well the individual difference of emotion ratings aligns with that of the neural signals, considering the inter-subject discrepancy across all situations.

**Fig. 2.**
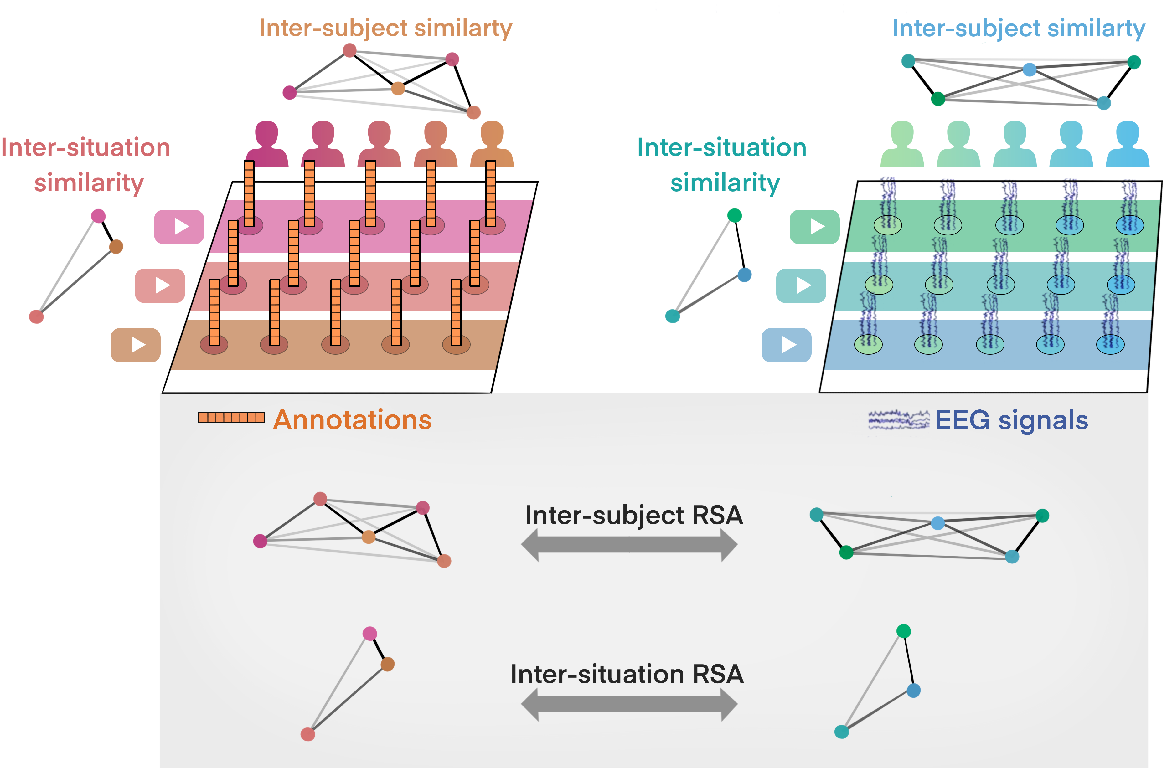
The overall illustration of inter-subject representational similarity analysis (RSA) and inter-situation RSA.

#### 2.4.1 Inter-situation RSA

In inter-situation RSA, for each participant, we generate a set of inter-situation similarity matrices based on their emotion annotations across all situations. Similarly, another set of inter-situation similarity matrices is generated using their neural signals across all situations^[11]^. The RSA process is conducted between the inter-situation similarity matrices of emotion ratings and the inter-situation similarity matrices of neural signals (Fig. 3a).

**Fig. 3.**
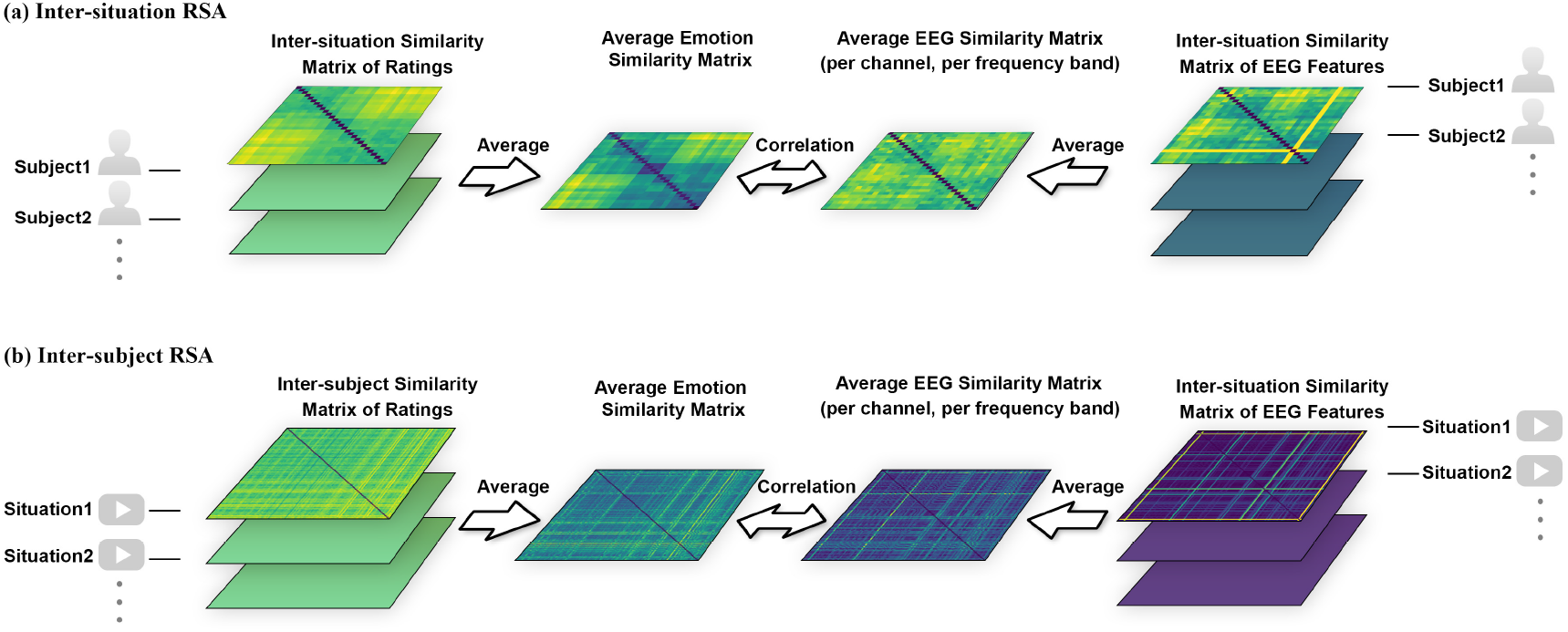
The pipeline of (a) inter-situation representational similarity analysis (RSA), and (b) inter-subject RSA.

(1) Inter-situation Similarity Matrix of Emotion Ratings: For each subject *i*, the inter-situation similarity matrix of emotion ratings 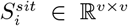 is computed based on all *E* emotion items, according to equation (3) and (4).

Inter-situation Similarity Matrix of Neural Signals: The Inter-situation similarity matrix of neural signals for participant *i*, 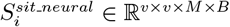 is computed from the EEG features of that participant in all *v*, situations, denoted as *X*_*i*_ ∈ ℝ^*v×M ×B*^, where *M* is the number of channels and B is the number of frequency bands.

Specifically, the signals are filtered into five frequency bands: Delta: 0.5–3 Hz, Theta: 4–7 Hz, Alpha: 8–13 Hz, Beta: 14–29 Hz, Gamma: 30–40 Hz . Next, frequency domain features are extracted, including Differential Entropy (DE) and Power Spectral Density (PSD). Difference between each feature dimension is computed to obtain the inter-situation dissimilarity matrix of neural signals 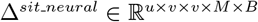.

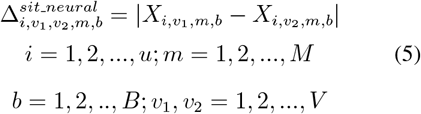

where 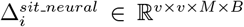 represents the intersituation difference patterns of participant *i*. Each element of it corresponds to the difference of an EEG feature between two situations. Then, the dissimilarity matrix Δ^*sit neural*^ was converted to similarity matrix *S*^*sit_neural*^ by normalizing it and subtracting it from an all-ones matrix.

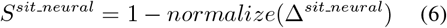

Here we compute the average inter-situation similarity matrix of ratings across all participants, obtaining 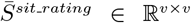, and the average inter-situation similarity matrix of EEG features for all participants, obtaining 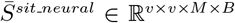. For each channel *m* and frequency band *b*, then compute the correlation coefficient between its neural similarity matrix and rating similarity matrix:

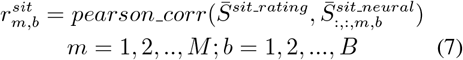

where *r*^*sit*^ ∈ ℝ^*M ×B*^ encompasses the inter-situation consistency between annotations and EEG across all channels and frequency bands.

#### 2.4.2 Inter-subject RSA

The computation of inter-subject RSA follows a structure similar to inter-situation RSA, with the key difference being that the similarity is calculated across subjects rather than situations. In inter-subject RSA, for a specific situation, a set of inter-subject similarity matrices is calculated based on the emotional annotations from all participants. Another set of inter-subject similarity matrices is generated using the neural signals from all subjects^[12]^. The correlation between these matrices is then used to assess the relationship between the inter-subject differences in emotion ratings and neural signals. (Fig. 3b). (1) Inter-Subject Similarity Matrix of Emotional Annotations: The inter-subject similarity matrix of situation *j* is denoted as 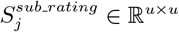, where *u* is the number of subjects. This is calculated based on the ratings of all subjects in this situation *R*_*j*_. When using multivariate emotion, *R*_*j*_ ∈ ℝ^*u×E*^ ; When using uni-variate emotion, 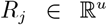; When using positive emotion, 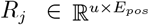; And when using negative emotion, 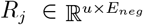. Next, 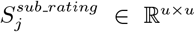 is obtained by first computing distance matrix 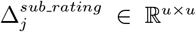 then normalize and substract it from a all-ones matrix, similar to the process shown in equation (3) and (4). 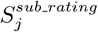 represents the inter-subject similarity pattern for situation *j. S*^*sub rating*^ ∈ ℝ^*v×u×u*^ was averaged over the situation dimension to obtain the similarity matrix of ratings 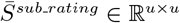. (2) Inter-Subject Similarity Matrix of Neural Signals: The inter-subject similarity matrix of neural signals across all situations, 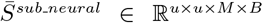, is obtained from the EEG features of all subjects in all situations, denoted as *X ∈* ℝ^*u×v×M ×B*^ . For situation *j* and subjects *i*_1_, *i*_2_, the computation is as follows:

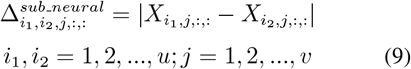

Here, 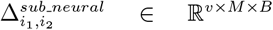 represents the neural dissimilarity between subjects *i*_1_, *i*_2_ calculated regarding *v* situations, *M* channels and *B* bands. Then, the dissimilarity matrix Δ^*sub neural*^ was converted into similarity matrix *S*^*sub neural*^ by normalizing it and subtracting it from an all-ones matrix.

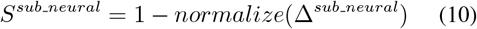

Then average *S*^*sub neural*^ ∈ ℝ^*u×u×v×M ×B*^ along the third dimension (situations), yielding the average neural similarity matrix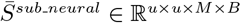.

The matrices 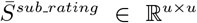 and 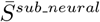 reflect the inter-subject similarity patterns across all situations in neural activity or ratings, i.e., inter-subject differences. Here, for each channel *m* and frequency band *b*, we calculate the correlation coefficient between its neural similarity matrix and the rating similarity matrix:

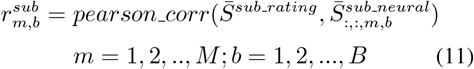

*r*^*sub*^ ∈ ℝ^*M ×B*^ represents the correlation of inter-subject ratings’ difference and EEG features’ difference.

## 3. Results

### 3.1 Inter-rater Reliability

For the inter-rater reliability test, ordinal representations showed significantly higher consistency between subjects than interval ratings (Fig. 4a), which is consistent with previous research.

**Fig. 4.**
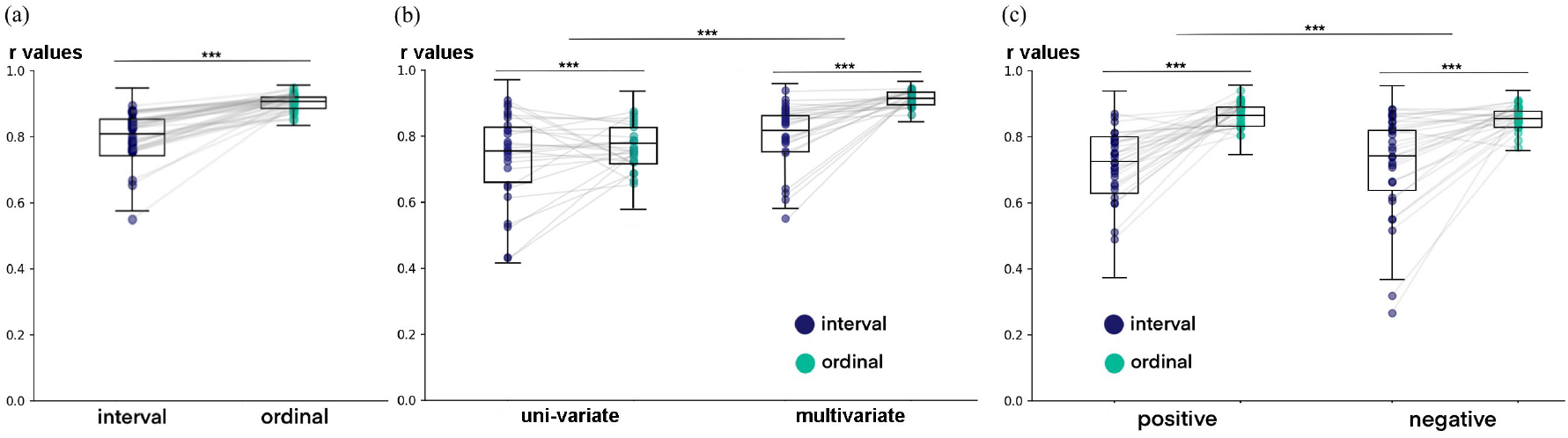
The results of reliability test. (a) The inter-rater reliability demonstrated by pairwise similarity of all subjects’ inter-situation similarity matrices using interval annotation and ordinal representation. (b) The inter-rater reliability of uni-variate and multivariate emotions. This is demonstrated by the pairwise similarity of inter-situation similarity matrices of interval and ordinal data, using univariate emotions and multivariate emotion. “Joy” is chosen as the uni-variate to be shown in the figure. Any other uni-variate emotions yielded lower inter-rater reliability than multivariate emotions (shown in Supplementary Figs. 1, 2). (c) The inter-rater reliability of positive and negative emotions. This is demonstrated by the pairwise similarity of inter-situation similarity matrices of positive (joy, tenderness, inspiration, amusement) and negative (anger, disgust, fear, sadness) emotions, using interval data and ordinal representation respectively.

When comparing multivariate and uni-variate emotions, ordinal representation resulted in a notable reliability improvement for both cases (Fig. 4b). On the other hand, multivariate emotion representations exhibit higher inter-subject reliability compared to uni-variate representations, regardless of which univariate emotion rating is used and whether ordinal representation is applied (Supplementary Figs. 1, 2). Using two-way ANOVA, we found a significant interaction between multivariate/uni-variate and ordinal/interval emotion representations (*ps <* 0.001 in statistical tests of all uni-variate emotions). This suggests that ordinal representation has much higher reliability than interval representations when combined with multivariate emotion.

When comparing positive and negative emotions, the ordinal representation improved the reliability of both (Fig. 4c, positive: *t*(7503) = 138.9, *p <* 0.001, negative: *t*(7503) = 104.6, *p <* 0.001), with a greater increase for positive emotions (two-way ANOVA: *F* (3, 30008) = 143.2, *p <* 0.001).

### 3.2 Validity and Neural Substrates of Ordinal Emotion Representations

The validity of emotion measurements is demonstrated by the maximum correlation coefficient between emotion annotation and EEG components. We visualized the correlation maps in inter-situation RSA and inter-subject RSA analyses (Fig. 5). We identified the maximum correlation coefficients among features across all channels and frequency bands and compared them between interval emotion representations and ordinal emotion representations. For both inter-situation RSA and inter-subject RSA, ordinal representations had a higher maximum correlation coefficient than their interval counterparts, indicating higher validity of ordinal representations. Regardless of the feature sets used (DE or PSD), or the type of representation (ordinal or interval), the brain regions most correlated with group-level emotional representations were concentrated in the temporal regions in higher frequency bands (beta and gamma). Moreover, the correlation in this region and frequency band was stronger under ordinal representation compared to interval representation. The ordinal representation brought validity improvements for both uni-variate and multivariate emotions.

**Fig. 5.**
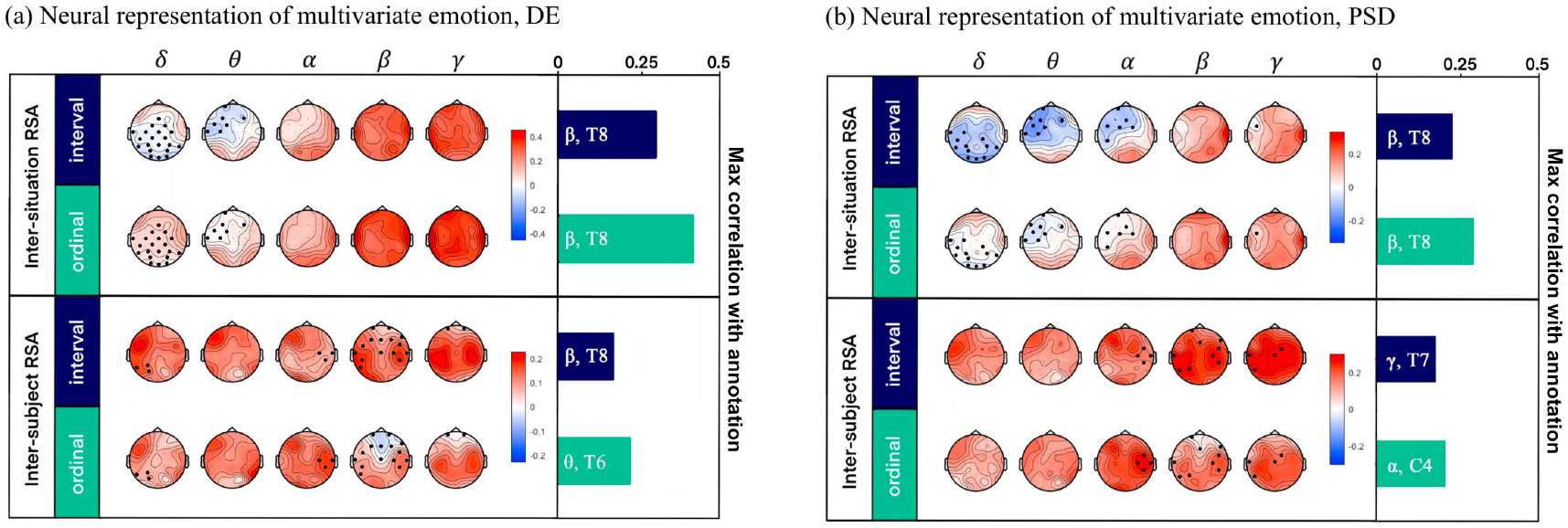
The neural representation of multivariate emotion (all items). The bars at right indicate the levels of validity, represented by maximum r values in RSA across EEG channels and frequency bands. The corresponding frequency bands and channels are shown in the bars. (a) Using differential entropy (DE) as the EEG feature. (b) Using power spectrum density (PSD) as the EEG feature. The channels with considerable differences (threshold = 0.1) between interval and ordinal are marked with black dots.

When comparing multivariate emotion representations with their uni-variate counterparts in RSA analyses, multivariate emotion representations consistently exhibit higher correlations with neural signals (Fig. 6). We further visualized the correlation patterns for inter-situation and inter-subject RSA across different uni-variate emotions. The inter-situation RSA between uni-variate ordinal ratings and EEG DE features showed a high correlation across widespread electrodes in beta and gamma bands, especially in temporal regions for positive emotions (Supplementary Fig. 3). These correlation maps align with the correlation maps for multivariate ordinal ratings in Fig. 5. For inter-subject RSA, the bilateral central, parietal and temporal regions in the gamma band had high correlations with most uni-variate emotion ratings (Supplementary Fig. 4). The left frontotemporal regions and right temporoparietal regions in delta and theta bands also showed correlation with uni-variate emotion ratings, especially for positive emotions. These patterns were also reflected in inter-subject RSA analyses of multivariate emotions (Fig. 5).

**Fig. 6.**
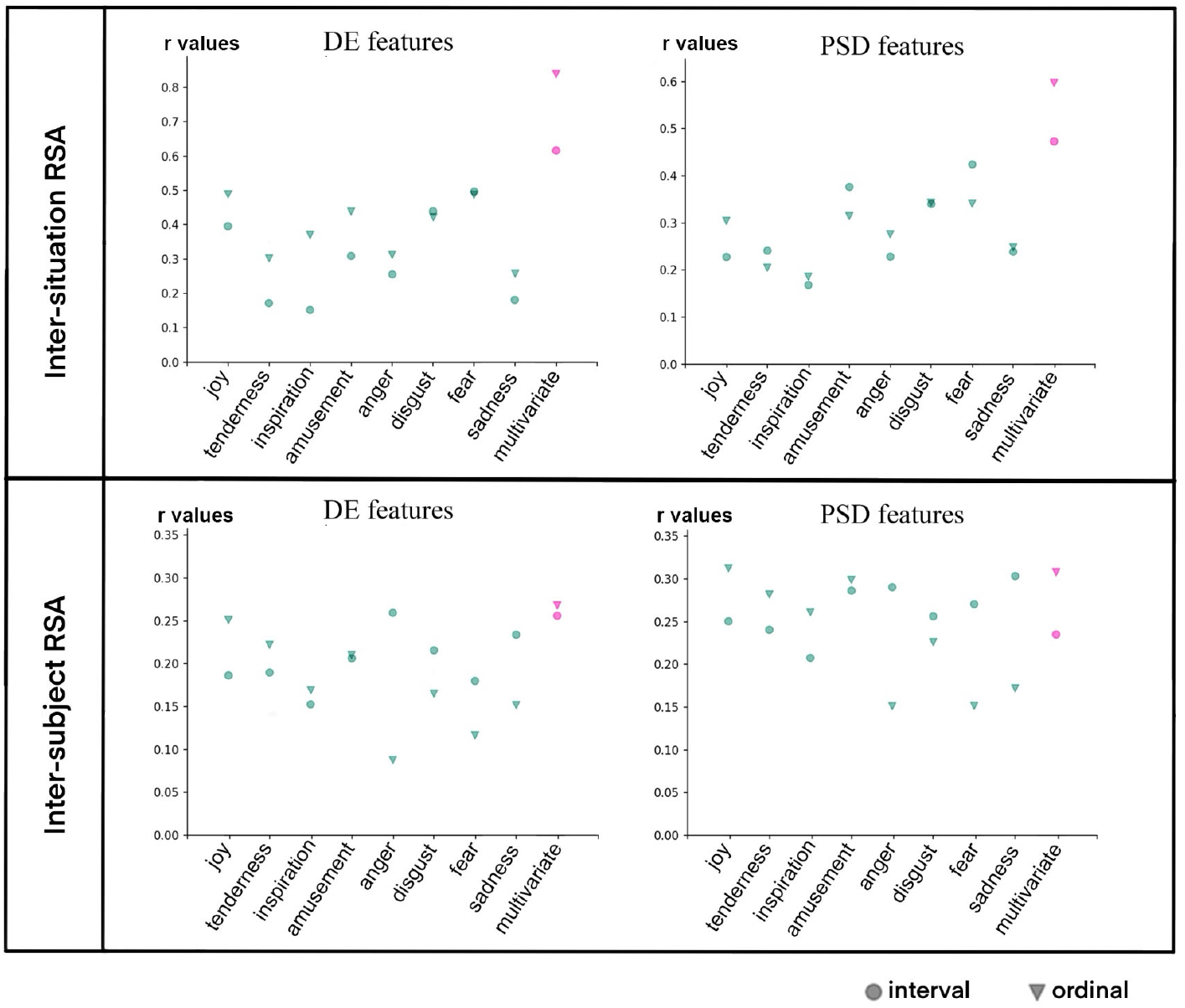
The validity demonstrated by maximum correlation coefficient between EEG signals and ratings, using uni-variate and multivariate emotion, and two different forms (interval form in circle, ordinal form in triangle). Above shows the result based on inter-situation RSA, demonstrating the validity in capturing group emotional commonalities. Below shows the result based on inter-subject RSA, demonstrating the validity in capturing individual emotional differences. The chosen EEG feature is differential entropy (DE) for left figures and power spectrum density (PSD) for right figures.

When comparing positive and negative emotions, ordinal representation significantly enhanced the validity of positive emotions, while its impact on negative emotions was inconsistent, showing both increases and decreases. Specifically, in inter-situation RSA, the validity of ordinal and interval data is comparable; however, in inter-subject RSA, interval data shows higher validity (Figs. 6, 7). Fig. 7 shows the validity described by the maximum correlation coefficient between emotion annotations and EEG components (DE features and PSD features), where positive emotions consistently show a greater increase when using the ordinal representation. Fig. 6 illustrates the validity of individual uni-variate emotion items under ordinal and interval conditions, where positive emotion items exhibit more stable and larger improvements. The EEG encoding of positive and negative emotions based on DE and PSD features are shown in Fig. 8 and Supplementary Figs. 5 and 6, respectively.

**Fig. 7.**
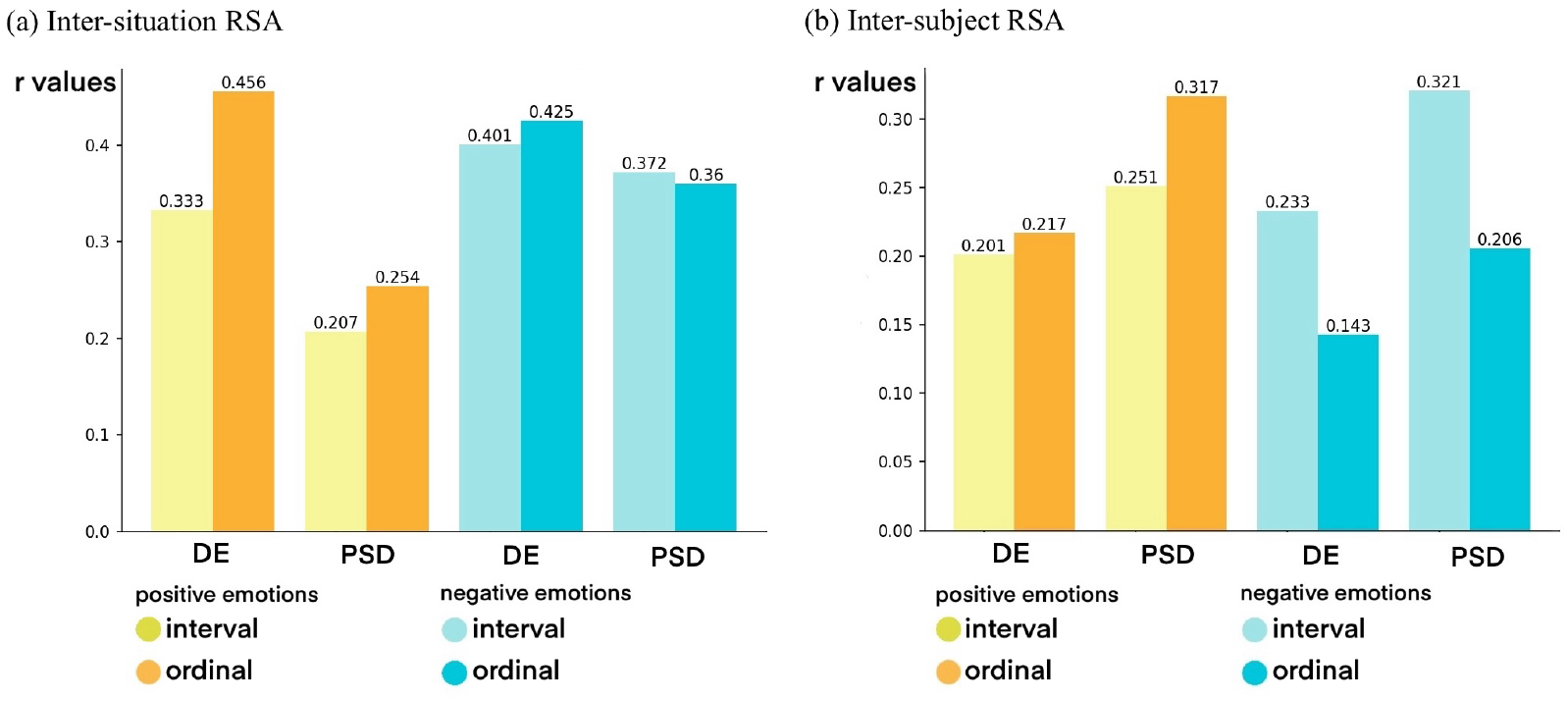
The validity demonstrated by maximum correlation coefficient between EEG signals and ratings, using positive and negative emotions, and two different forms. Ordinal representation brings greater improvements for positive emotions than negative emotions. (a) validity in inter-situation RSA (b) validity in inter-subject RSA. EEG features including differential entropy (DE) and power spectrum density (PSD) are used, respectively.

**Fig. 8.**
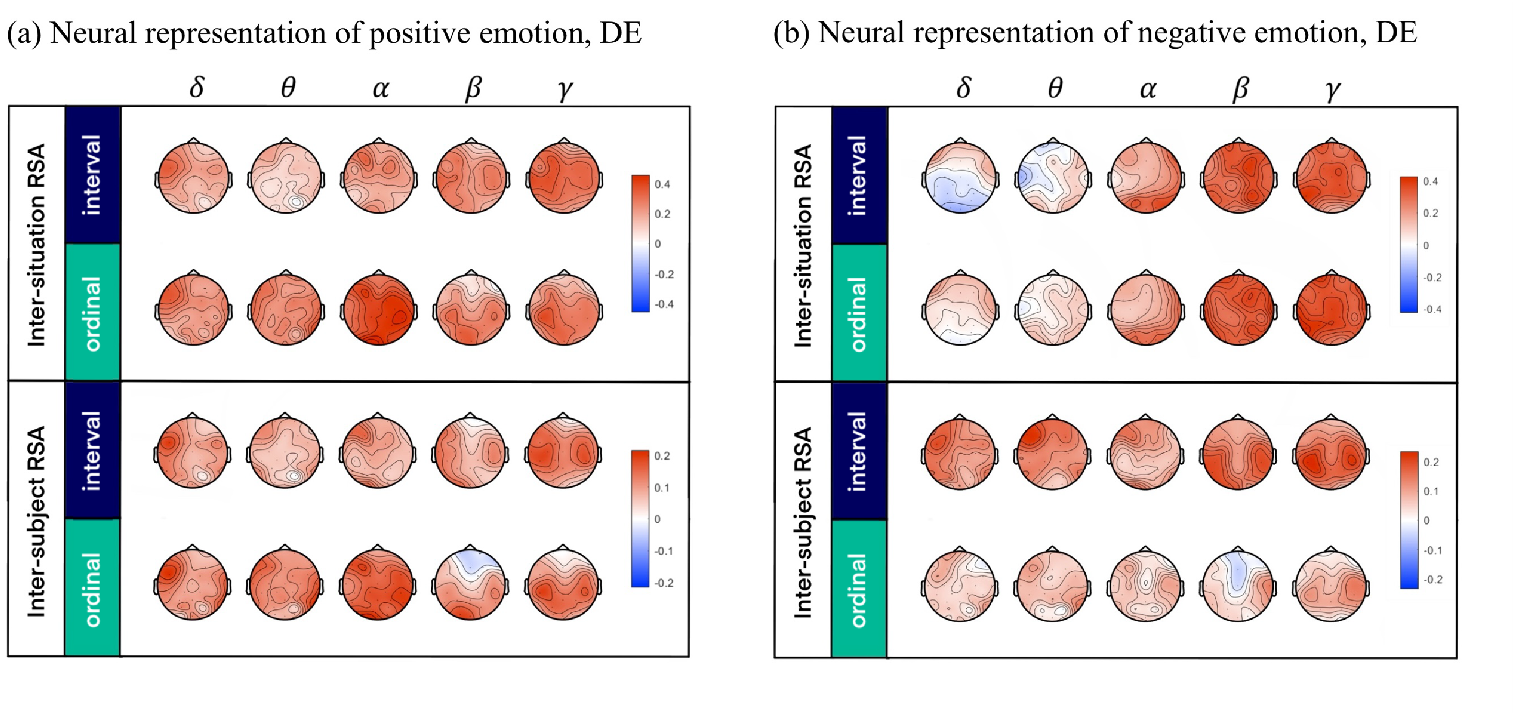
The neural representation of positive emotions (joy, tenderness, inspiration, amusement) and negative emotions (anger, disgust, fear, sadness). (a) Positive emotions. (b) Negative emotions. Shown in the figure are the results based on differential entropy (DE) features. For results based on power spectrum density (PSD) features, see supplementary materials.

## 4 Discussion

In this study, we used a large-scale emotional EEG dataset to uncover the neural correlates of ordinal emotional representations. Our findings revealed that ordinal representations not only exhibit greater interrater reliability (consistent with previous studies^[3,19]^), but also better capture commonalities and individual differences in the characteristics of EEG. Moreover, multivariate emotions correspond more closely to neural signals compared to each uni-variate emotion.

The main contribution of this work is the first use of EEG data to demonstrate the validity of multivariate ordinal emotions and reveal their neural substrates.

The noise in emotional self-reports has long been a topic of discussion. In most emotional datasets, self-reported emotions are recorded as vectors in interval form. This form of measurement has been shown to lack reliability when significant changes in emotions occur^[3,5]^. However, such dynamic changes in emotions are the key focus of research in affective computing and affective science. The lack of robustness arises from the noise introduced during the externalization of emotions, where participants map the intensity of emotional experiences onto an interval space. Yannakakis et al. summarized extensive theoretical and experimental evidence, demonstrating that this noise stems from the mismatch between the intrinsic nature of emotional experience and the interval scale^[3]^. In relative judgment models, context serves as an anchor that helps define the scale for assessing an option’s relative position. Individuals evaluate items in relative terms, rather than absolutely, demonstrating a strong reliance to anchors^[20]^. In the process of emotion self-report, the brain lacks an absolute scale to evaluate emotional intensity and instead relies on comparisons between temporally adjacent emotional instances. The relative relationships between emotional instances do not conform to Euclidean space, and embedding them into Euclidean space during externalization introduces distortions. There are numerous alternative Euclidean outputs that can equally satisfy the same relative relationships. When considering multiple subjects and trials, the arbitrary differences among these alternatives introduce additional noise. For example, the same score given by two participants may not represent the same emotional experience.

Although the scores are noisy, the relative relationships should remain stable. Compared to interval measurements, ordinal measurements focus only on the relative ranks between data points, without considering distances or ratios between them. Therefore, using ordinal measurements for emotional experiences should largely eliminate the noise present in interval-based emotional self-reports. Studies have demonstrated that ordinal measurement can reduce noise in emotional self-reports by examining the inter-subject reliability of emotion ratings and their correspondence with peripheral physiological signals or emotional speech features^[3,8–10, 19]^. This study further demonstrated the validity of ordinal ratings with EEG signals and revealed the neural substrates of ordinal emotion representations. Specifically, EEG features in temporal regions, particularly in high-frequency bands such as beta and gamma, showed a stronger correlation with ordinal emotion representations. This is consistent with previous research suggesting that these brain regions and frequency bands play a critical role in emotion processing^[21–23]^.

To benchmark against existing emotion measurement methods, we further evaluated the performance of ordinal and categorical (i.e., nominal) approaches using the RSA pipeline. In measurement theory, ordinal measurement conveys an amount of information that lies between categorical (nominal) and interval measurement^[3]^. Specifically, categorical measurements provide the least structural information (e.g., mere classification without order), while ordinal measurements introduce additional information through rank ordering. In this study, for the categorical data, we represent emotional state of a subject in a specific trial by the emotion item with the highest score, thereby reducing the emotional experience to a single category. Our results show that the validity of categorical measurements is comparable to that of ordinal measurements (Supplementary Fig. 7). This finding indicates that reducing the information provided by ordinal scales does not enhance data quality but rather imposes significant limitations due to information loss. The drawbacks of categorical measurements have been well documented, primarily because human emotions are too complex to be described by a single category^[24, 25]^. Therefore, compared to interval and categorical methods, the ordinal approach is preferable, as it combines high reliability and strong neurophysiological correspondence with the capacity to capture multidimensional emotional states.

One compelling discovery in this work was the superior alignment of multivariate ordinal emotion representations with EEG signals, compared to their uni-variate counterparts (Fig. 6). This implies that considering the complexity and co-occurrence of multiple emotional states provides a more robust measure of emotional experiences. This finding aligns with the understanding that human emotions are rarely experienced in isolation but rather as complex, intertwined states^[26, 27]^. A more detailed investigation of the correlation maps for uni-variate emotion ratings revealed that activities in temporal regions, particularly in high-frequency bands such as beta and gamma exhibited high correlations with multiple emotions (Supplementary Figs. 3, 4). These correlation patterns also aligned with those observed in multivariate analyses, suggesting that the EEG representations of different uni-variate emotions partially overlap. Multivariate emotion ratings provide a richer and more nuanced representation of emotional states, resulting in stronger correlations with neural signals (Fig. 6).

Another interesting finding is that ordinal representation consistently improved the reliability and validity of positive emotions, whereas its impact on negative emotions was less stable. This suggests that the externalization patterns of the selected positive emotions in this dataset (amusement, inspiration, joy, tenderness) are more aligned with ordinal structures than the selected negative emotions (anger, fear, disgust, sadness). This may be because the four positive emotions are more akin to social emotions^[28]^, which are thought to involve more cognitively sophisticated processes, including self-reflection and attribution^[28]^. These emotions have complex co-occurrence patterns^[29–31]^ and are highly context-dependent^[32]^. Such complexity could make them more difficult to be evaluated by an internal interval scale^[33]^ and largely rely on comparisons, exhibiting a more pronounced ordinal structure. In contrast, most negative emotions in this dataset are considered to lack the involvement of higher-order cognitive processes and are experienced more directly^[32]^. These emotions might exhibit less noise during the externalization process.

These findings provide new insights into how we conceptualize and measure emotions. Viewing emotions through an ordinal representation offers not only more reliable emotion annotations but also hints at underlying neural mechanisms of emotion processing. Specifically, emotion processing in the human brain may involve divisive normalization, a computational mechanism where a neuron’s activity is modulated by the overall activity of its surrounding context^[3]^. This mechanism introduces context dependence, as the value of a stimulus is encoded relative to other available stimuli. Commonly associated with processes like sensory adaptation and attention^[34]^, divisive normalization may similarly play a role in the relative encoding of emotions, as suggested by the evidence in this study. Ordinal emotion representations, in turn, align well with this mechanism by reflecting context-dependent evaluations.

In terms of applications, our findings have multifaceted implications for affective computing. First, they offer a framework for designing scoring paradigms and improving the reliability of emotion annotations through ordinal scales^[11]^. Second, they provide a foundation for developing emotion decoding algorithms; for instance, Yannakakis et al. proposed contrastive learning methods inspired by ordinal annotation schemes^[35]^. Third, ordinal emotion representations serve as a denoising tool for emotion self-reports, facilitating the study of cross-individual, cross-group, and cross-cultural differences in emotional experiences. These applications collectively address significant limitations in current affective computing practices, particularly the challenges of noise and variability in emotion self-reports, paving the way for more robust and scalable approaches.

It should be noted that this work still has several limitations. First, we used only a single dataset. The robustness of the ordinal representation’s EEG encodings across different datasets and experimental paradigms remains to be studied. Second, whether the characteristics of ordinal representation exhibit robustness across cultures and demographics is also a topic that requires investigation. Third, whether more cognitively sophisticated emotions, such as social emotions, indeed exhibit a more pronounced ordinal structure is another direction worth further exploration.

## 5 Conclusion

Our study demonstrates that ordinal emotion representations offer a more effective approach to capturing the complexity of human emotions. Compared to interval-based measurements, ordinal representations show higher inter-rater reliability and better alignment with both group commonalities and individual differences in neural activity. Notably, ordinal representations enhance the alignment with neural signals particularly when applied to multivariate and positive emotions. This highlights the co-occurrence of multiple emotions and intricate patterns of positive emotions are better reflected by ordinal scales. In summary, ordinal representations provide a robust framework for measuring emotions, ensuring greater inter-rater reliability and stronger grounding in neurophysiological data. These findings are essential for accurately representing human emotions in machines, improving machines’ capacity to comprehend human emotions.

## Acknowledgment

The authors wish to thank Zongsheng Li for his valuable suggestions. This work was funded by Shenzhen Science and Technology Innovation Committee (RCBS20231211090748082, 2022410129, KCXFZ20201221173400001, KJZD20230923115221044), the National Natural Science Foundation of China (62472206), Shenzhen Excellent Youth Project (RCYX20231211090405003), and SUSTech Undergraduate Innovation and Entrepreneurship Training Program (2024S07).

---

**Xuyang Chen** received the B.S. degree in bioinformatics from Southern University of Science and Technology, Shenzhen, China, in 2024. He is working as a research assistant in the Department of Biomedical Engineering, Southern University of Science and Technology. His current research interests focus on the neural mechanisms of emotion evaluation and emotion experiences.

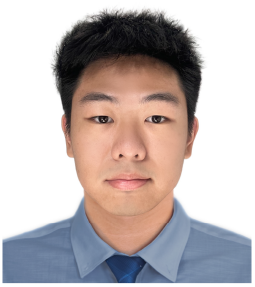

**Xin Xu** is an undergraduate student in the Department of Biomedical Engineering at Southern University of Science and Technology, enrolled in 2022. She is interested in affective computing, brain-computer interfaces and emotion-related neural representations.

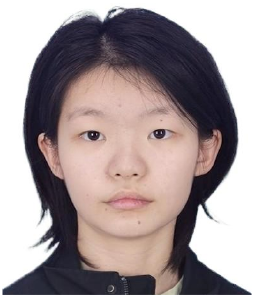

**Dan Zhang** received the B.E. degree in automation in 2005 and the Ph.D. degree in biomedical engineering in 2011, both from Tsinghua University, Beijing, China. He was a postdoctoral fellow in School of Medicine, Tsinghua University from 2011 to 2013. He is currently an associate professor at the Department of Psychological and Cognitive Sciences, Tsinghua University, Beijing, China. His research interests include social neuroscience, engineering psychology, and brain-computer interfaces. He is a member of the IEEE.

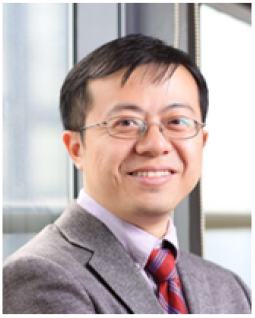

**Xinke Shen** received the B.E. degree in biomedical engineering from Beihang University, Beijing, China, in 2017 and the Ph.D. degree in biomedical engineering in Tsinghua University, Beijing, China, in 2023. He is currently a postdoctoral fellow in the Department of Biomedical engineering, Southern University of Science and Technology, Shenzhen, China. His research interests include affective computing and affective neuroscience.

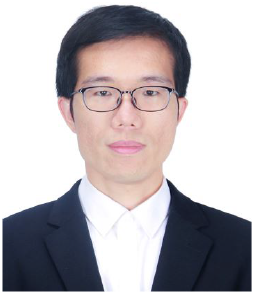

