## Supplementary material for "Exploring the Neural Basis and Validity of Ordinal Emotion Representation through EEG"

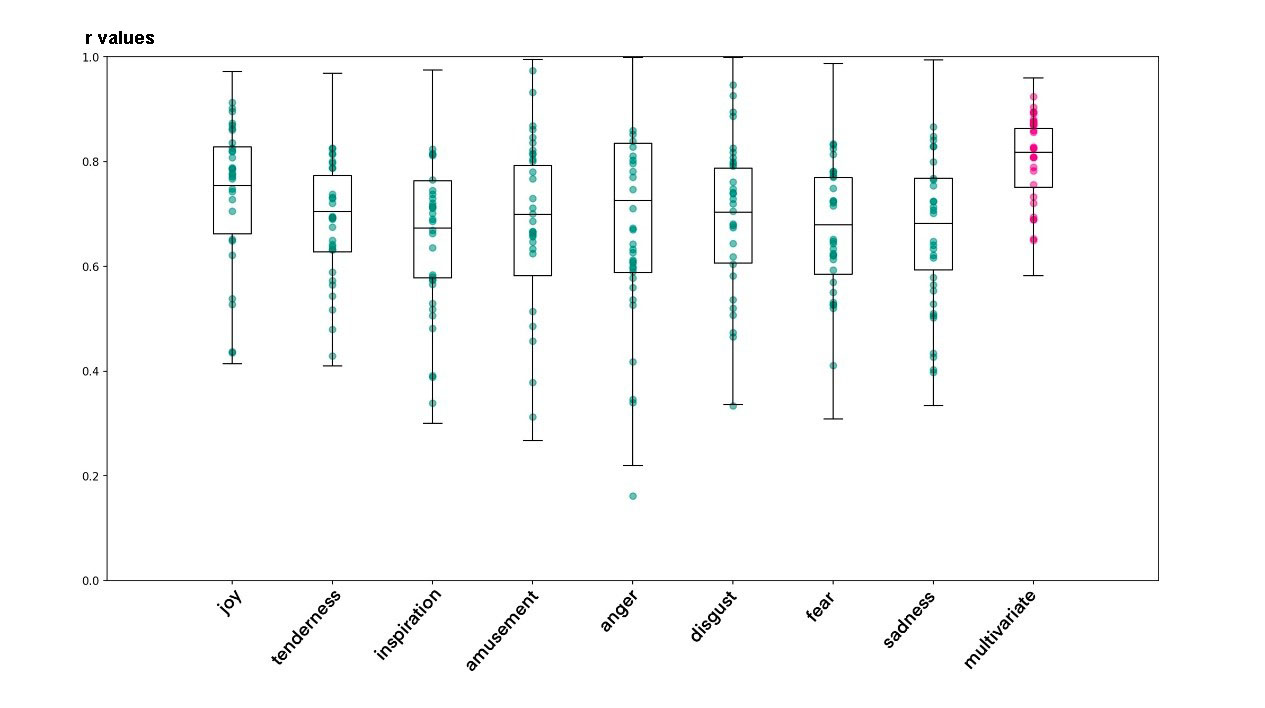

Supplementary Fig. 1. The inter-rater reliability of uni-variate and multivariate emotions, based on interval ratings. This is demonstrated by the pairwise similarity of inter-situation similarity matrices of the interval data, using uni-variate emotions and multivariate emotion. The differences between multivariate emotion and all uni-variate emotions are significant (p<0.001).

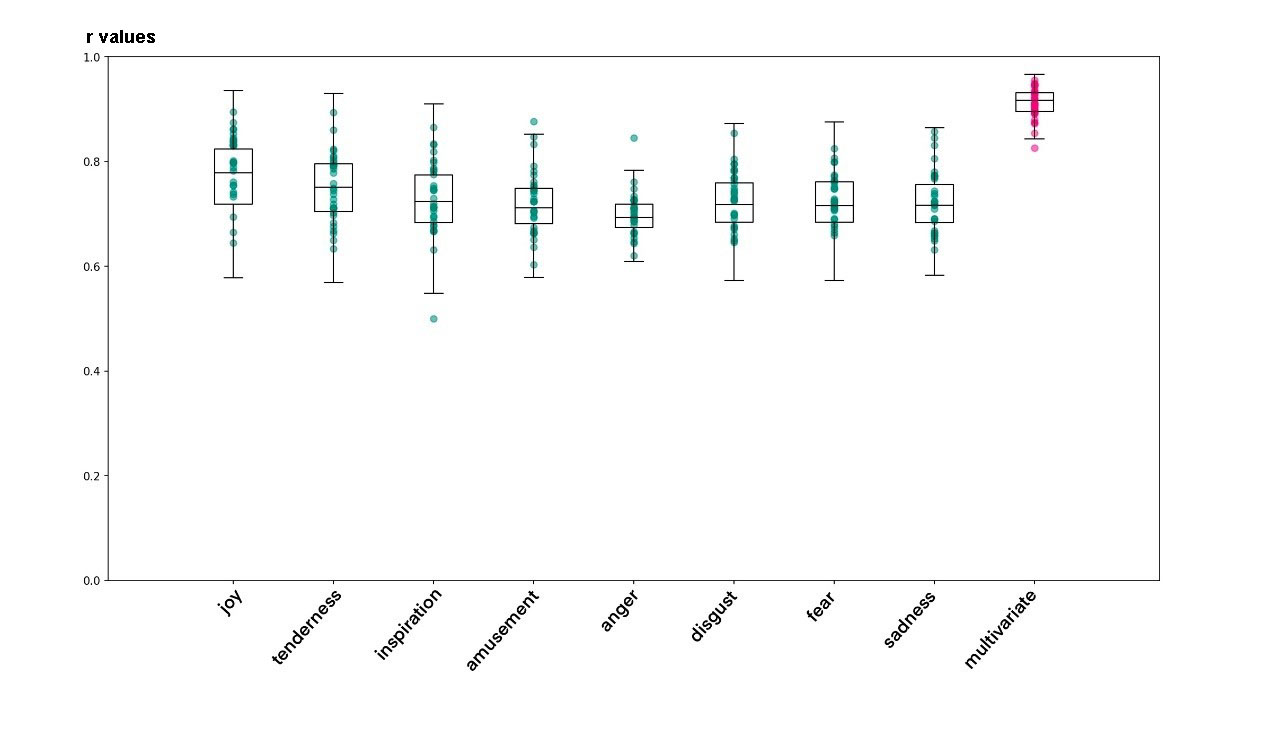

Supplementary Fig. 2. The inter-rater reliability of uni-variate and multivariate emotions, based on ordinal representation. This is demonstrated by the pairwise similarity of inter-situation similarity matrices of the ordinal data, using uni-variate emotions and multivariate emotion. The differences between multivariate emotion and all uni-variate emotions are significant (p<0.001).

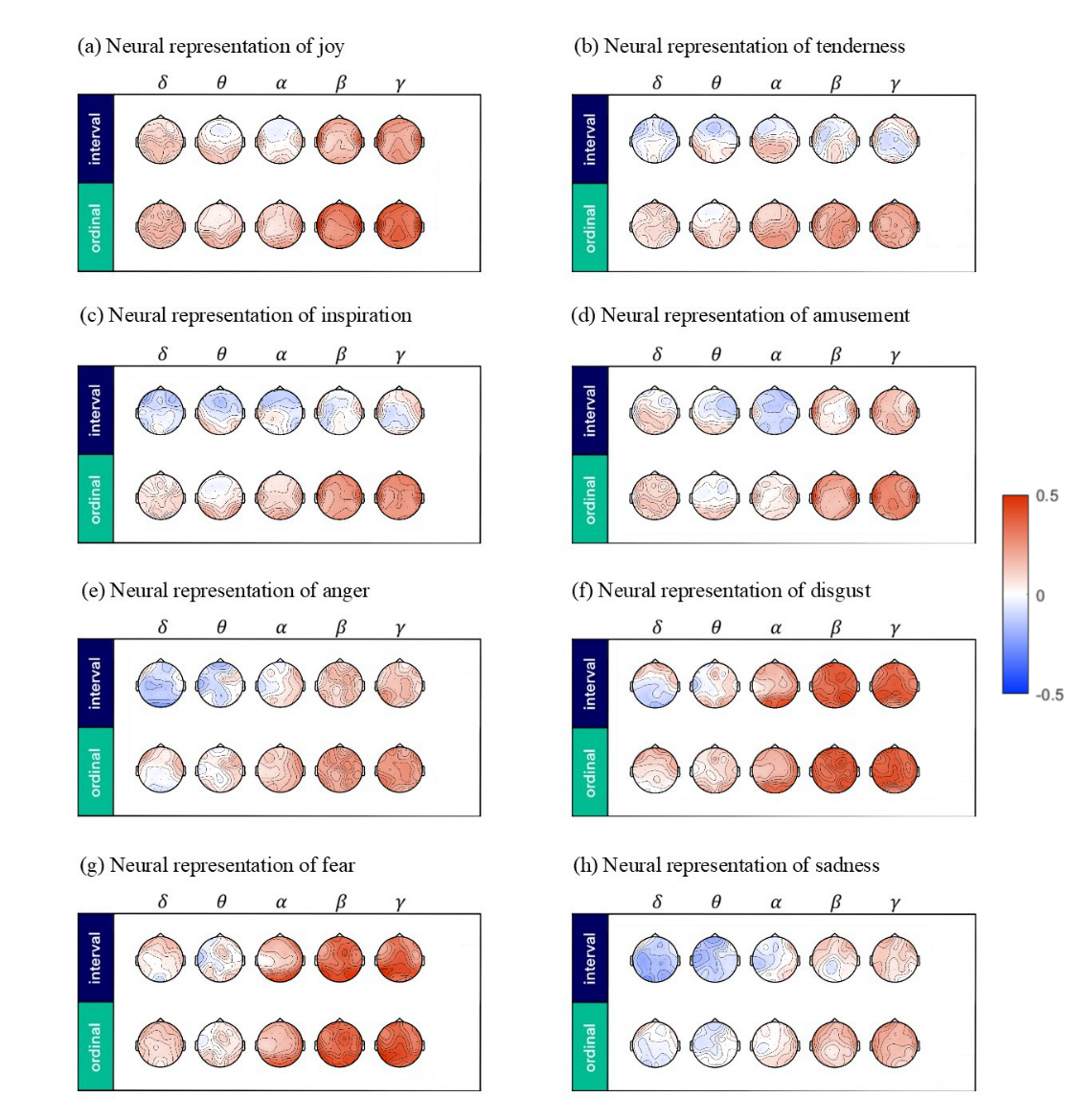

Supplementary Fig. 3. The neural representation of uni-variate emotions based on inter-situation RSA and differential entropy (DE) features.

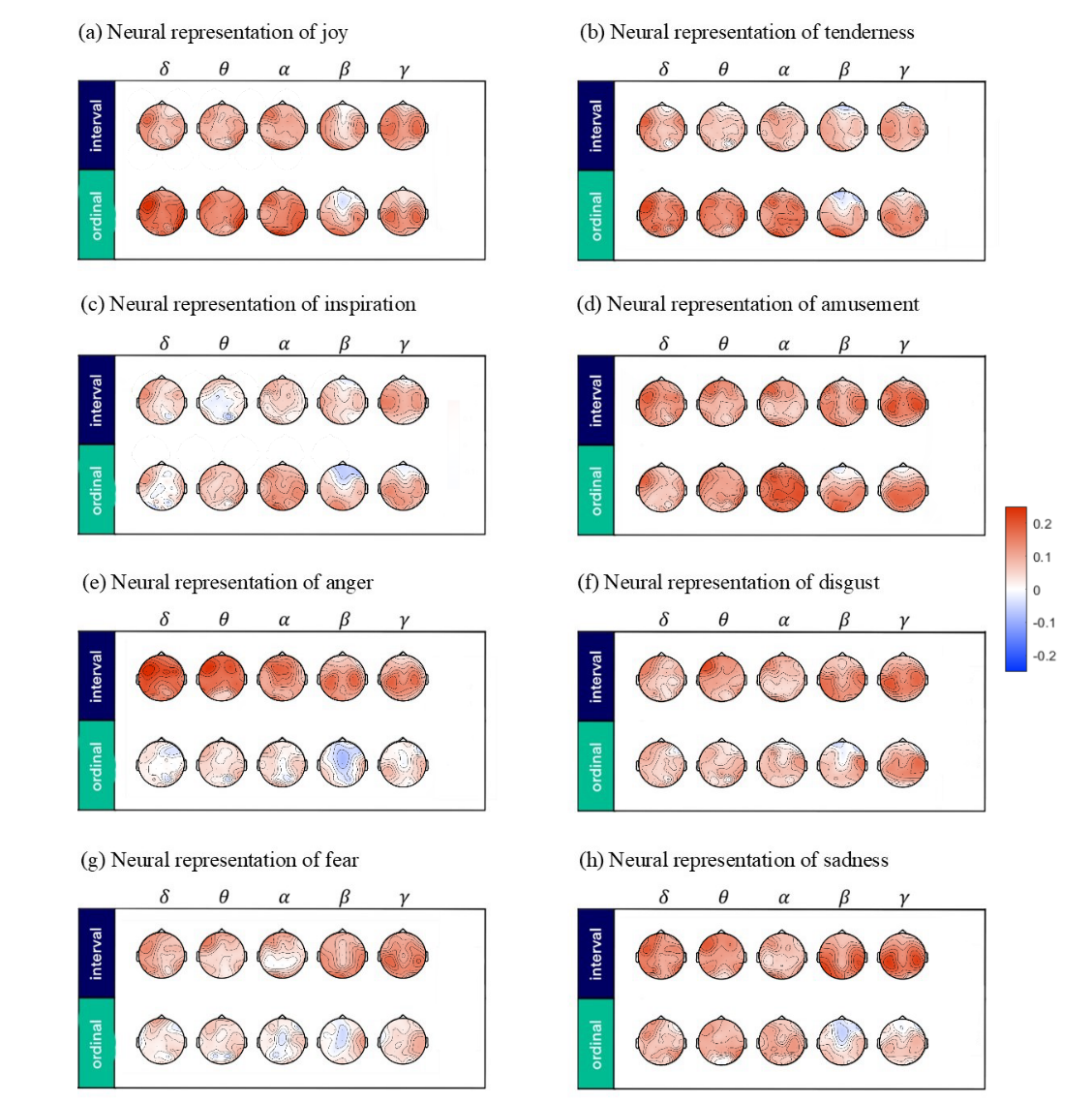

Supplementary Fig. 4. The neural representation of uni-variate emotions based on inter-subject RSA and differential entropy (DE) features.

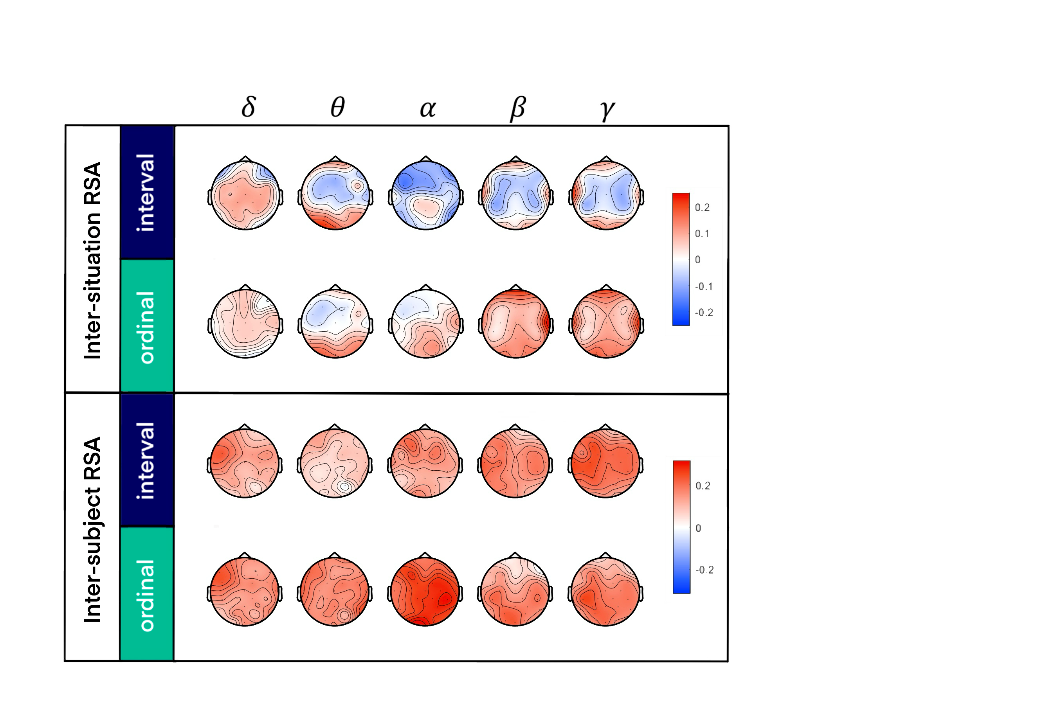

Supplementary Fig. 5. The neural representation of positive emotions (joy, tenderness, inspiration, amusement), based on power spectrum density (PSD) features.

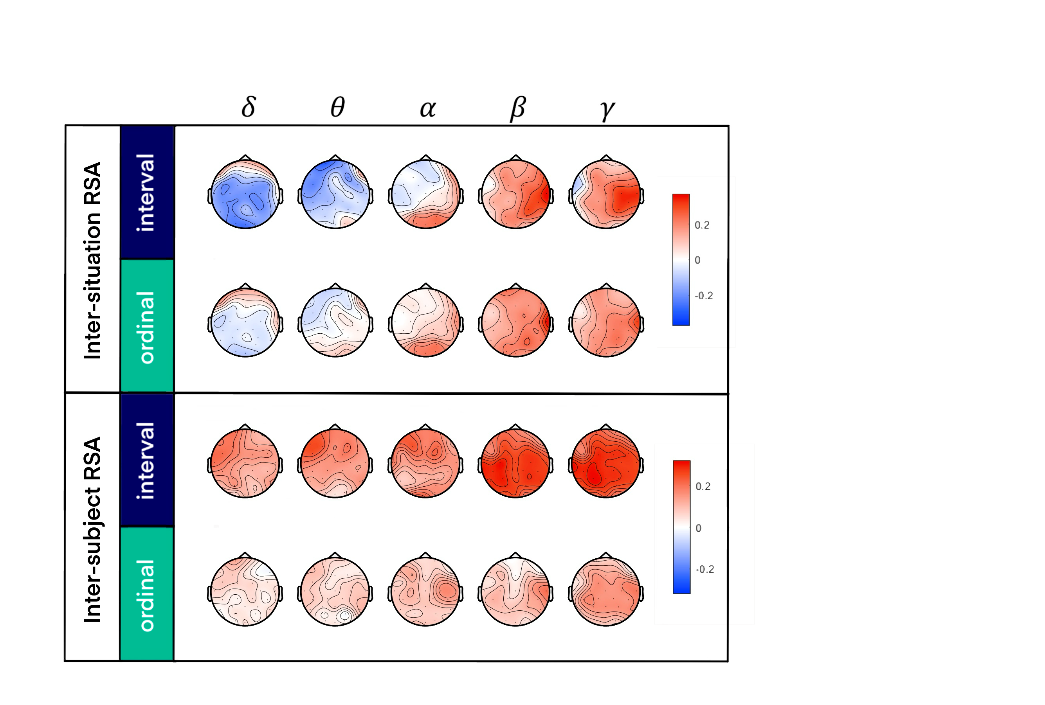

Supplementary Fig. 6. The neural representation of negative emotions (anger, disgust, fear, sadness), based on power spectrum density (PSD) features.

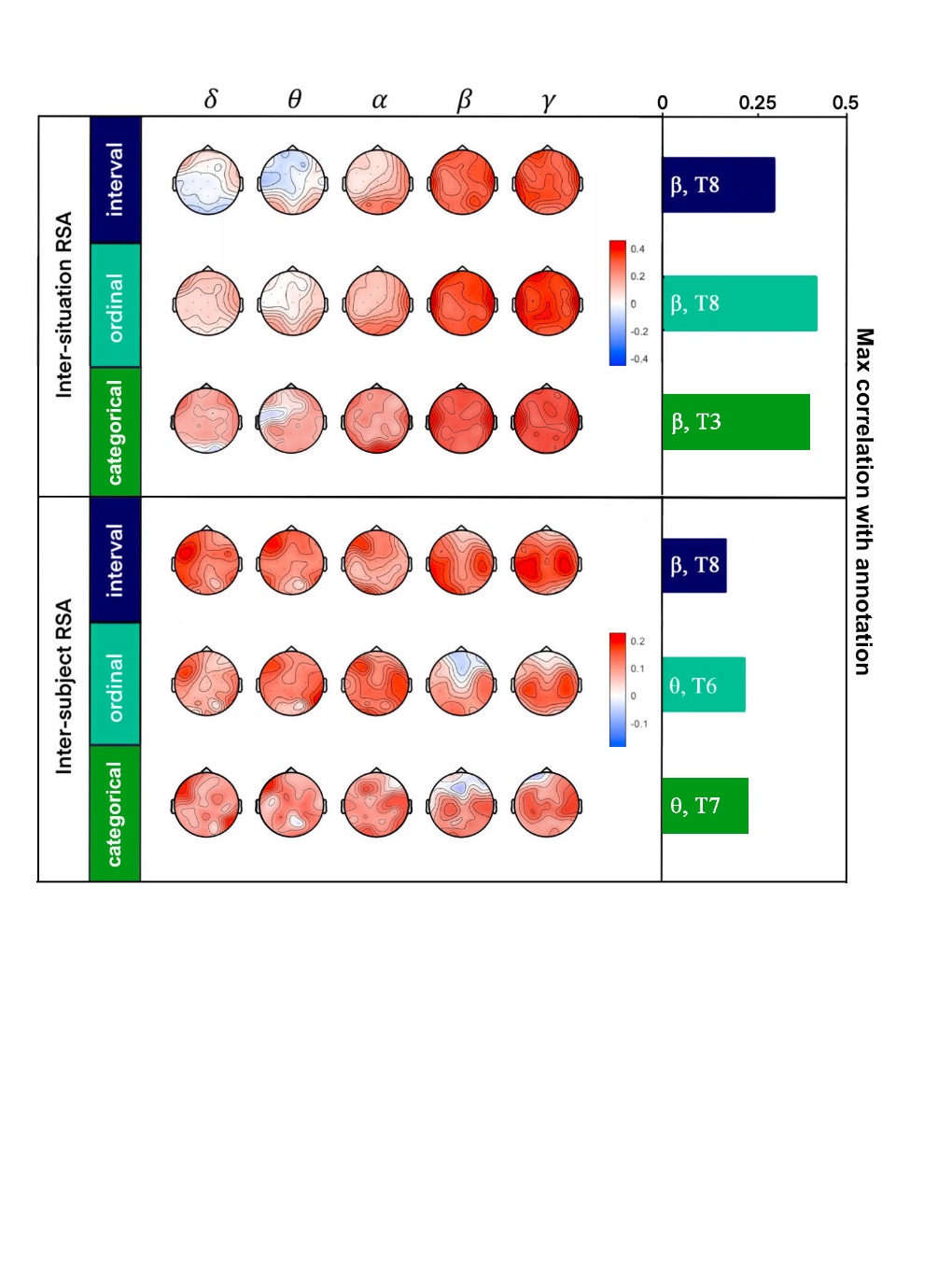

Supplementary Fig. 7. The neural representation of multivariate emotion, using interval, ordinal, and categorical measurement, respectively. The result is based on differential entropy features.
